# Balance control is more affected in older than younger adults by repeated visual perturbations during walking

**DOI:** 10.64898/2026.05.08.723731

**Authors:** Yaqi Li, Eugénie Lambrecht, Sjoerd M. Bruijn, Jaap H. van Dieën

## Abstract

Sensory degradation with aging can impair balance control, partly by disrupting visual contributions to self-motion estimation. We investigated age-related differences in center of mass (CoM) trajectories controlling in the frontal plane during exposure to repeated visual perturbations in walking. We hypothesized that the older adults would show larger responses to visual perturbations and less adaptation to repeated visual perturbation exposure. We applied three visual perturbations to 14 healthy older (age: 75.0±2.4) and 16 younger adults (age: 23.4±3.9) walking on a treadmill: fixating a stationary target with the background moving to the right (MB), tracking a target moving rightward over a stationary background with head rotation (MT-HR), and tracking a moving target with eye movement only (MT-EM). CoM position deviations due to the visual perturbations were assessed. Over the whole trial, the older adults did not show larger CoM position variability than the younger during visual perturbation walking, which was contrary to our hypothesis. During visual perturbation epochs, both age groups deviated in the same direction except MB. In MB, the older adults deviated to an opposite direction after a few perturbation repetitions. Moreover, in MT-HR and MT-EM, the older adults deviated earlier than the younger adults and they deviated more in the MT-HR condition. This indicates that older adults exhibit reduced ability to accurately estimate self-motion through correction by other sensory modalities when exposed to visual perturbations. Over repeated perturbations, the older adults showed decreased CoM deviations in MT-EM, which suggests that they still maintain the capacity to downweight visual information after repeated exposure.

## Introduction

As bipedal animals, humans face a substantial challenge to balance their center of mass (CoM) over a relatively small base of support [1]. To achieve this during walking, the main mechanism is to select an appropriate location for foot placement in relation to the ongoing movement of the CoM [2]. To this end, the CoM state is estimated by the integrating sensory information from visual, vestibular, proprioceptive, and cutaneous input [3–5].

The degeneration of sensory systems in older adults is one of the reasons for an age-related decline in balance control [6]. In older adults, impaired vestibular function causes a reduction in sensitivity to changes in spatial orientation [7] and impaired balance control [6]. Thus, older adults may rely more on the visual system for balance control [8–10]. However, visual information may provide erroneous information about body orientation in specific situations.

When encountering objects of interest, it is typical to fixate or track them using eye or head movements. Fixating on a stationary target while the background is moving is known to perturb balance control [11]. Such background movement can lead to an illusion of self-motion in the direction opposite to the background movement [12, 13]. For instance, when an individual is exposed to a moving visual background, they may perceive self-motion or postural sway despite remaining stationary. Visual tracking of moving targets could be another potential visual perturbation during walking. Visual tracking will generate a relative movement of the visual background on the retina. It may therefore induce a similar illusion as described above. Eye movement and head rotation during visual tracking may lead to changes in vestibular and proprioceptive information as well as mechanical effects on balance. All these different visual challenges occur constantly in daily life, but, despite their relevancy, the resulting balance responses and age-related differences are currently unknown. In our previous study [14], we exposed younger adults to three visual perturbation conditions during walking. We found that when participants visually fixated on a stationary target while the background was moving, their foot and CoM trajectories deviated in the direction opposite to the background movement. When participants visually tracked a horizontally moving target with head rotation or eye movement while the background was stationary, their CoM trajectories deviated in the direction of the target, but this happened several seconds after the target had started moving and after it had reached its stationary position. Given the higher visual reliance reported in older adults, such visual perturbations may generate greater effects in this group, and, as such, may pose greater challenges to balance control.

Increased visual dependence of balance control in older adults may be reflected in two ways. Firstly, because balance control ability declines with aging, older adults may be more susceptible to interference from sensory perturbations. Consequently, older adults may respond to visual perturbations earlier and stronger than younger adults. For example, the older adults may display earlier and larger CoM deviations during a target movement. Secondly, older adults may be less able to correct the resulting deviations by using other sensory information such as vestibular and proprioceptive information. Consequently, older adults may have more persistent responses to visual perturbation during walking. Also, older adults may show impaired sensory reweighting in a changing environment [15]. Generally, when environmental conditions change, reweighting of the visual, vestibular and somatosensory information can help to maintain balance [16]. It’s critical to with unpredicted environment changings. With aging, the adaptation to a new environment may be slower [17–19]. Thus, older adults may experience larger CoM deviations to a first visual perturbation and be less able to adapt during repeated exposure to visual perturbations.

We conducted a treadmill walking experiment in which we exposed older adults to three types of visual perturbations; 1) fixation on an object while the background moved, and visual tracking of an object moving over a stationary background by 2) head rotation or 3) eye movement only. We compared the results with the data obtained in younger adults from our previous study. We hypothesized that compared to younger adults, in all visual perturbation conditions: (1) variability of the CoM trajectory over the whole walking trial will be greater in older adults; (2) during visual perturbations, greater peak CoM deviation will occur in older adults; (3) older adults will show decreased adaptation to repeated visual perturbation exposure.

## Methods

### Participants

We recruited 20 healthy older and 18 healthy younger participants. Six older participants were not able to finish all the tests and their data were discarded. Data from two younger participants were discarded because of technical issues. Hence data from 14 older (5 female; 9 male) and 16 younger (10 female; 6 male) participants were used in this study. All of the participants reported normal visual acuity with or without glasses correction (older adults: 2 without glasses, 12 with glasses; younger adults: 8 without glasses, 8 with glasses). All participants were self-reportedly free from any neurological or musculoskeletal disorders (i.e. vestibular disorders, stroke, Parkinson disease etc.) and were not taking any medication known to negatively affect their balance or walking performance. All participants walked without any assistive devices. None of them had fallen in the previous three months. General information including height, body mass and date of birth was collected. Ethical approval was obtained from the ethics committee of the Faculty of Behavioral and Movement Sciences at Vrije Universiteit Amsterdam (VCWE-2023-170). All procedures conformed to the Declaration of Helsinki and participants signed informed consent before participation [20].

Three clinical scales were used to define participants’ cognitive and ambulatory status before the experiment. Cognitive status was tested by The Montreal Cognitive Assessment (MoCA) (“© MoCA Test Inc. All rights reserved.), which was designed as a rapid screening instrument for mild cognitive dysfunction. The maximum score is 30 points; a score of 26 or above is considered normal. Ambulatory status was defined by the Tinetti Performance Oriented Mobility Assessment (POMA) (Reprinted with permission, Mary E. Tinetti, M.D.© Copyright; 2006) and Dynamic Gait Index (DGI) [21]. POMA is a test which measures balance and gait ability. The assessment has a maximum score of 28 points. A total score between 25 and 28 points indicates a low risk of falling, a score between 19 and 24 points suggests a moderate risk, and a score below 19 points reflects a high risk of falling. DGI also assesses the likelihood of falling in older adults. The full score is 24 points, those who score below 20 points face high risk of falling, those who score more than 22 points are safe ambulators.

### Instruments

All measurements were conducted on a system that integrated a motion capture system sampling at 50 samples/sec (Optotrak, Northern Digital, Waterloo ON, Canada) with an instrumented treadmill (Motek-Forcelink, Amsterdam Netherlands), and a projector. Clusters of three markers were affixed to the participant’s bodies at the following locations: the posterior surfaces of both heels, the sacrum at the level of S2, the trunk at the level of T6, and the back of the head. A screen (2.50 m wide, 1.56 m high) was placed 1.60 m in front of the participants with the projector behind it, to present the visual perturbations. **Figure 1A** shows the full set-up.

**Figure 1.**
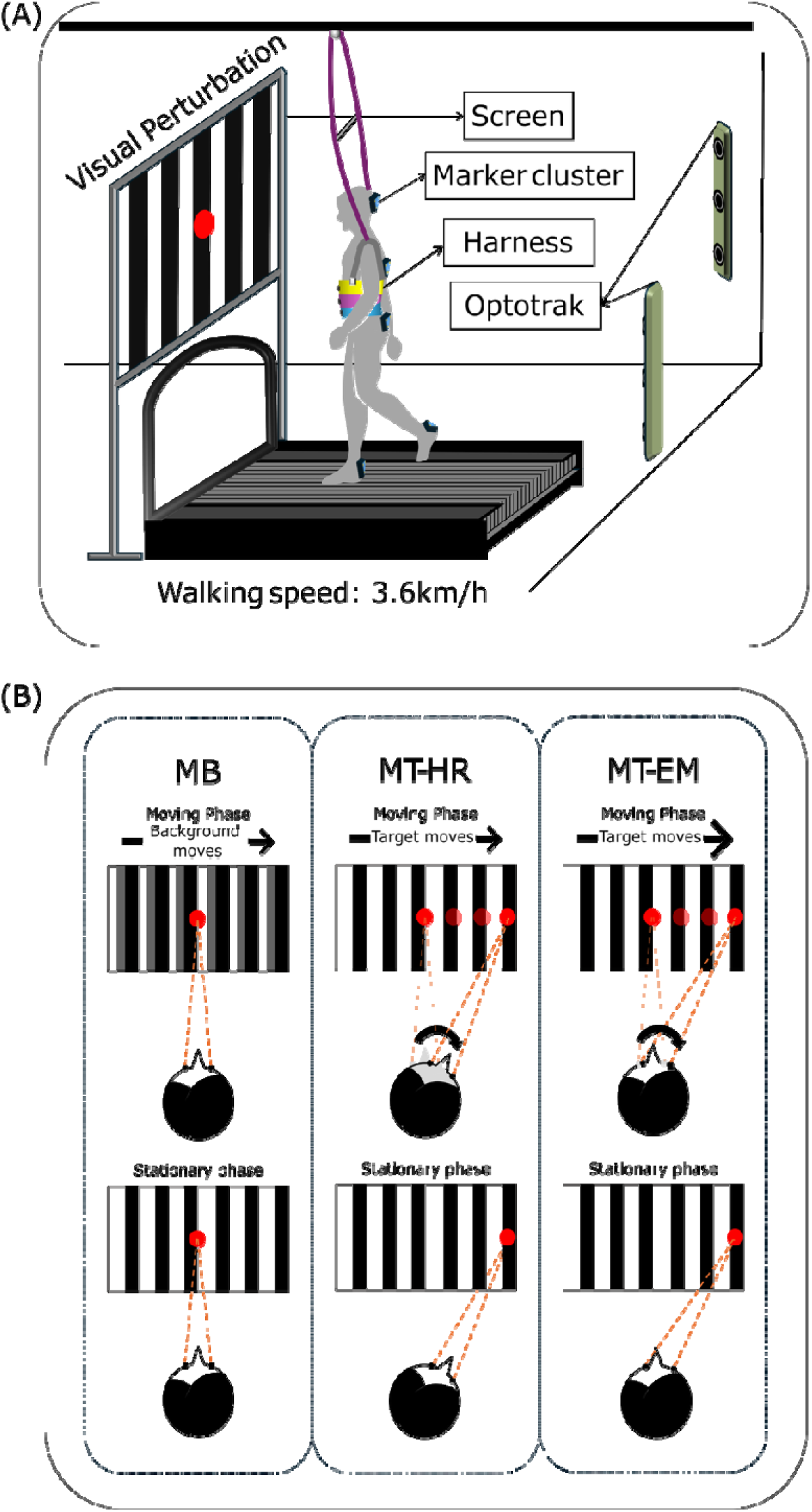
Illustration of the experimental setup (A) and visual perturbation conditions (B)

### Visual perturbations

A background of 12 uniformly distributed black-and-white vertical stripes (0.205 m * 1.560 m) was presented on the screen. A red target dot measuring 12 cm in diameter was positioned at the center of the screen. The height of the red target was adjusted to be at the participant’s eye level. Participants walked under four different conditions: (a) Normal walking (NW), in which the whole scene was stationary and participants were asked to walk normally. (b) Moving background (MB) perturbations, during these perturbations, the red target dot was fixed in the middle, while the black-and-white background moved horizontally from the middle to the right side corresponding to 45 degrees in 4s, stayed at the side for 8s, then went back to the middle. In this condition, participants were asked to keep looking at the stationary red target dot in the middle. (c) Moving target with head rotation (MT-HR), during these perturbations, the black-and-white background was stationary, while the red target dot moved horizontally to 45 degrees from the center to the right side in 4s and stayed at the side for 8s then moved back to the middle. Participants were asked to track the moving target. (d) Moving target with eye movement (MT-EM), during these perturbations, the scene was the same as during MT-HR. Participants were asked to track the target using their eyes only, while keeping their head stationary. All conditions are illustrated in **Figure 1B**.

Visual perturbations were triggered at right heel strikes. In each walking trial, we provided approximately 14 perturbations in total based on feasibility ascertained during pilot testing. The first perturbation was triggered at the 20^th^ right heel strike. Six to eight heel strikes occurred randomly after every perturbation before a new perturbation was provided.

### Procedures

Data from the younger adults in this study were previously reported [14]. All data were collected in a single study and hence all procedures were identical between groups, except that a ceiling-mounted harness was only used for the older participants to guarantee their safety. The treadmill speed was set to 3.6 km/h. To enhance immersion, the lights in the lab were turned off, and the curtains were closed. Before the actual measurement, there was a 10-minute familiarization period at the same walking speed and with all perturbations. Subsequently, a normal walking trial was conducted, followed by the three perturbation trials presented in a random order. Each trial lasted 5 minutes. A 5-minute seated rest was given between trials.

### Data analysis

The mediolateral CoM position was approximated as the average position of the pelvis marker cluster. Mediolateral foot positions were represented by both heel clusters. Heel strikes were determined by the maximum anteroposterior position of the heel marker.

Gait characteristics reflecting frontal plane balance were calculated over the whole walking trial, including step width, step width variability, step frequency, and CoM position variability. Step width was defined as the mediolateral distance between the heel positions of both feet at heel strike. Step frequency was defined as the number of steps per minute. Step width variability was quantified as the standard deviation over steps. CoM position variability was quantified as the standard deviation averaged over phases of the time-normalized stride. We analysed the time series of foot position and CoM position in detail from the start of the visual perturbation to its end. We divided these episodes into two phases: moving and stationary phases (**Figure 1B**). The continuous variables were low-pass filtered at 0.25Hz (2nd order bidirectional Butterworth filter) to eliminate the fluctuations related to the stride cycle. To facilitate comparison between trials, we subtracted the value at the first sample of the visual perturbation epoch from all subsequent measurements. Peak position during each phase was selected as the maximum absolute value. We determined the average values of these variables among participants over perturbations.

### Statistics

All statistical analyses were performed in Matlab (R2023a, The Mathworks Inc., Natick, MA). For all statistical tests, the assumption of normality was checked using a Shapiro-Wilks test. For all repeated measures analyses, if the assumption of sphericity was violated, the Greenhouse-Geisser correction was be applied to adjust the degrees of freedom. For all tests, the significance level (*p*-value) was set at 0.05. We calculated the effect size to provide an estimate of the magnitude of observed effect: *d* (Cohen’s d) for one-sample t-tests and independent t-test and 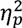 (Partial eta squared) for ANOVA.

To assess baseline comparability between older and younger groups, we performed independent sample t-tests on height, weight and three clinical scales. For the variables MoCA and DGI which did not follow the normal distribution, Mann-Whitney U tests were used instead.

To assess the first hypothesis that the older adults would show increased CoM position variability during visual perturbation, we conducted repeated measures analyses of variance with two factors (Visual perturbation and Age). We used a Box-Cox transformation on CoM position variability which did not follow a normal distribution. If we found main effects or interaction effects to be significant, we performed post-hoc comparisons with Bonferroni correction to check the differences between age groups and perturbation conditions.

To test the second hypothesis that the older adults would show larger peak CoM deviations during the visual perturbations, we did two tests. First, we performed one-sample t-tests to assess if averaged peak CoM position was significantly different from zero. Then, we performed repeated measures analyses of variance with three factors (Visual perturbation, Phase, Age, as well as their interactions) for peak CoM position in the perturbation epochs (MB, MT-HR and MT-EM). If we found significant interaction effects among the three factors, we performed independent t-tests in each perturbation condition and phase to test the differences per age group. If not, then we tested the interaction effects of two factors (Visual perturbation and Age). A post-hoc test was subsequently performed to assess the differences between age groups in each perturbation condition.

To test the third hypothesis that the older adults would show less adaptation to repeated visual perturbations, we fitted linear mixed effects models to quantify the change in peak absolute CoM deviation over repeated perturbations. We fitted the model with the data in the first 13 visual perturbation epochs (the minimum perturbation numbers across all walking trials) for each perturbation condition. The model included fixed effects Repetition (from 1 to 13), Age (older adults 0; younger adults 1) and the Repetition X Age interaction. Random intercepts for each participant accounted for repeated measures within subjects.

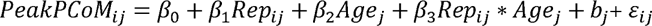

where *i* denotes the repetition number, and *j* denotes group number (older adults = 0, younger adults = 1). *PeakPCoM_ij_* represents the peak absolute CoM position measured at repetition *i* for group *j*. *β*_0_ is the intercept. *β*_1_ represents the fixed effect of repetition (*Rep_ij_*). *β*_2_ represents the fixed effect of age group (*Age_j_*). *β*_3_ represents the interaction effect between repetition and age group. *b_j_* represents as subject-specific random intercept. *ε_ij_* is the residual error for each repetition per group.

### Sample-size justification

We conducted power analysis for three-way repeated measures ANOVA with Age, Phase and Visual perturbation. We assumed a moderate effect size (f = 0.25), a significance level of *α* = 0.05, and a statistical power of 80% (1 − *β* = 0.80). Power analysis results indicated a total sample size of 30 participants would be sufficient to detect the expected effects.

For the linear mixed model, we performed a Monte Carlo power analysis based on a moderate expected interaction effect between Age and Repetition. The simulation included two groups and repeated measurements across 13 times points. For each total sample size, 500 datasets were independently simulated and analyzed using the same linear mixed model as that used in the current analysis. The results indicated that a total sample size of 30 participants provided 92% power to detect the expected interaction effect.

## Results

### Participants demographics

Demographics and data on the cognitive and physical condition of the two groups of participants are listed in **Table 1**. There were no significant differences between older and younger adults in height, weight and MoCA score. The result of MoCA and POMA tests showed both older and younger adults had good cognitive status and good balance and gait ability. Compared with the younger adults, DGI scores were significantly decreased in the older adults.

**Table 1.** Demographics and data on the cognitive and physical condition of older and younger participants and the statistic results for the group comparison. MoCA: The Montreal Cognitive Assessment; POMA: Tinetti Performance Oriented Mobility Assessment (POMA); Dynamic Gait Index (DGI).

|  | Old (n = 14) | Young (n = 16) | pValue |
| --- | --- | --- | --- |
| Age (yr) | 75.0±2.4 | 23.4±3.9 | - |
| Height (m) | 1.72±0.11 | 1.70±0.09 | 0.499 |
| Weight (kg) | 71.9±12.0 | 62.2±11.7 | 0.135 |
| MoCA | 26.6±2.2 | 27.9±1.1 | 0.07 |
| POMA | 28±0 | 28±0 | - |
| DGI | 22.6±1.2 | 23.7±0.7 | **0.003** |

Note: Significant differences are presented in bold font.

### Effects of visual perturbations over the whole trial

**Figure 2** shows the CoM position variability over the whole trial in the two age groups. There was no significant Visual perturbation X Age interaction effect on CoM position variability (*p* < 0.412, 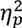 = 0.033). But there were significant effects of Visual perturbation (*p* < .001, 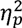 = 0.638) and Age (*p* < .001, 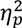 = 0.452). Post-hoc comparisons showed that CoM position variability in all perturbation conditions was significantly increased compared to NW (MB: *p* <.001, *d* = 1.499; MT-HR: *p* <.001, *d* = 2.476; MT-EM: *p* < 0.001, *d* = 2.259). The older adults showed larger CoM position variability than the younger adults (*p* < 0.001, *d* = 1.155).

**Figure 2.**
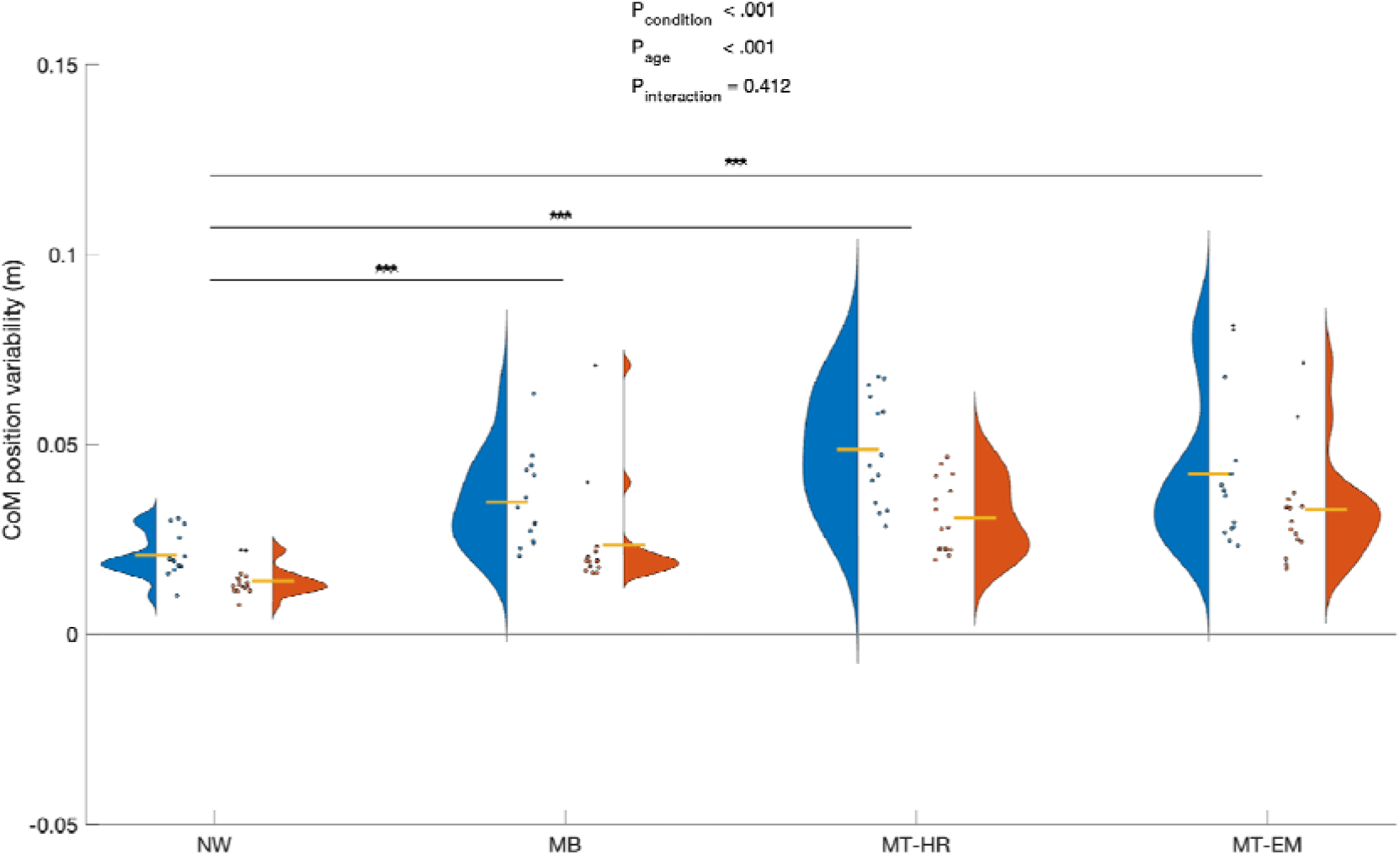
Effects of Visual perturbation conditions on CoM position variability in older and younger adults. The yellow lines represent for the mean values of each group. NW: normal walking, MB: moving background, MT-HR: moving target with head rotation, MT-EM: moving target with eye movement. Significant between groups are indicated by an asterisk on the top

### Destabilizing effects during the visual perturbation

Qualitatively, CoM and foot trajectories changed during the visual perturbations (**Figure 3**). In the MB condition, the younger participants consistently deviated in the direction opposite to the background movement (left, negative), while the older participants did not deviate consistently during the visual perturbation epoch. Inter-individual variance between the older participants was large. In the MT-HR condition, both older and younger participants deviated in the direction of the target (right, positive). Compared with the younger participants, who started to deviate at the end of the moving phase, the older participants deviated earlier, starting already at the beginning of the moving phase, and continued to deviate throughout the stationary phase. The deviation over the whole visual perturbation epoch was larger in the older than in the younger participants. In the MT-EM condition, both older and younger participants deviated in the direction of the target. However, the older participants started their deviation at the beginning of the moving phase, earlier than the younger participants. They both continued to deviate throughout the stationary phase. The between-participant variability was large relative to the magnitude of the effects throughout the perturbations.

**Figure 3.**
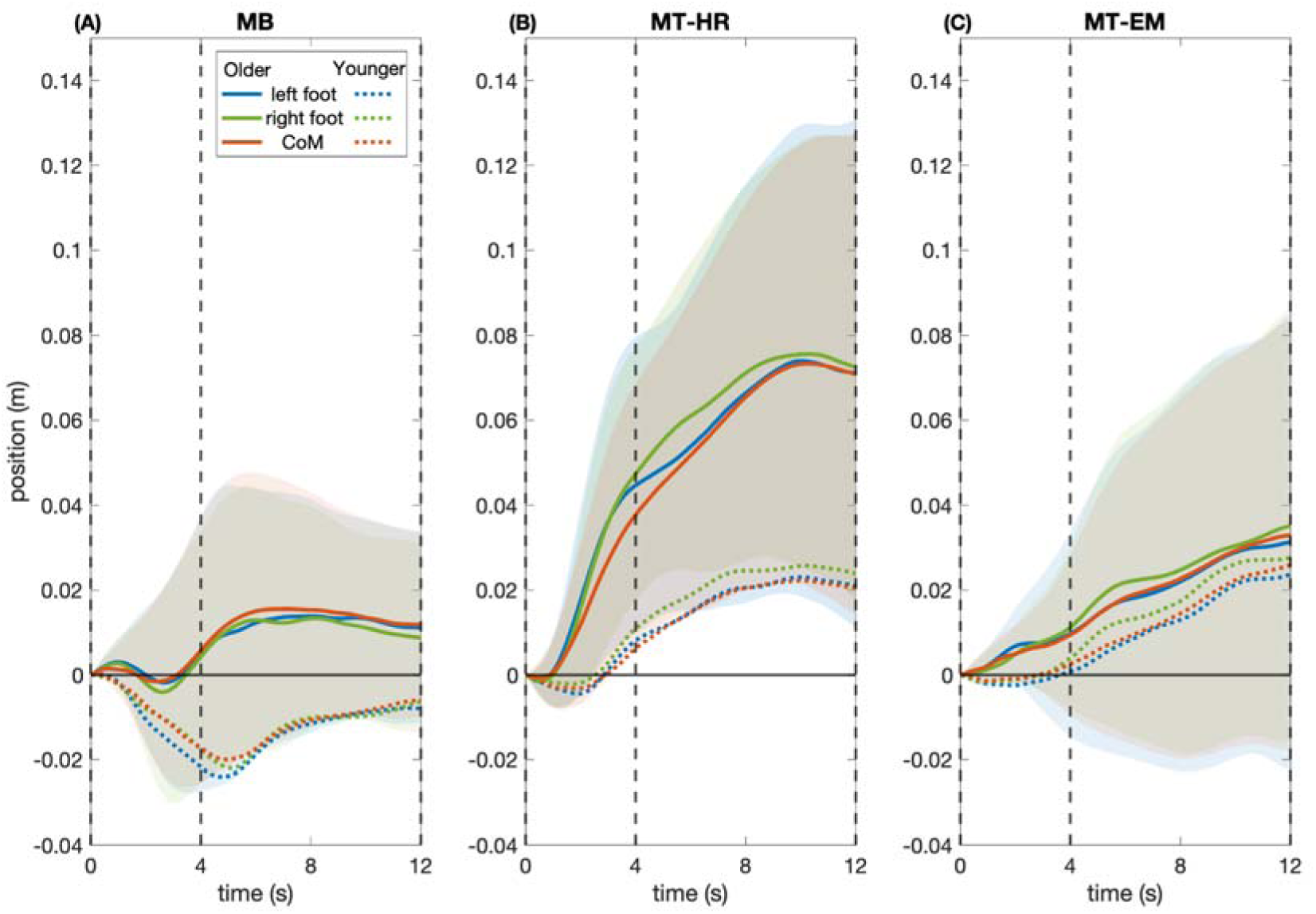
Deviations in CoM and foot positions during visual perturbation epochs in older (n = 14, mean± s.d.) and younger adults (n = 16, mean± s.d.). All the data were referenced to the first sample of the visual perturbation epoch. The shaded area represents the between-participants standard deviations only for the older adults. For the variance in the younger adults, omitted here for readability, we refer to Li, Lambrecht et al. [14]. From left to right: (A) MB: moving background. (B) MT-HR: moving target with head rotation, (C) MT-EM: moving target with eye movement. The vertical dashed lines separate the different phases: the line at 0s marks the start of the moving phase, the line at 4s marks the start of the stationary phase, and the line at 12s marks the end of the stationary phase

Statistical analysis of the peak CoM positions relative to the start of the visual perturbation (**Figure 4**, **Table 2**) confirmed the qualitative descriptions above. In the MB, in the older participants, the averaged peak CoM positions were not significantly different from zero neither in the moving nor in the stationary phase (*p* = 0.682, *d =* 0.112; *p* = 0.129, *d =* 0.433), while, in the younger participants, significant effects were found in both phases (moving phase: *p* <.001, *d =* -1.604; stationary phase: *p* <.001, *d =* -2.135). In the MT-HR and MT-EM conditions, in older adults, CoM position was significantly larger than zero in both the moving and stationary phases (MT-HR: *p* = 0.004, *d =* 0.949; *p* < .001, *d =* 1.621; MT-EM: *p* = 0.032, *d =* 0.641; *p* = 0.039, *d =* 0.613) indicating deviations in the moving target direction. However, younger adults only showed significant deviations during the stationary phases of both MT-HR (*p* = 0.026, *d =* 0.619) and MT-EM conditions (*p* = 0.041, *d =* 0.559).

**Figure 4.**
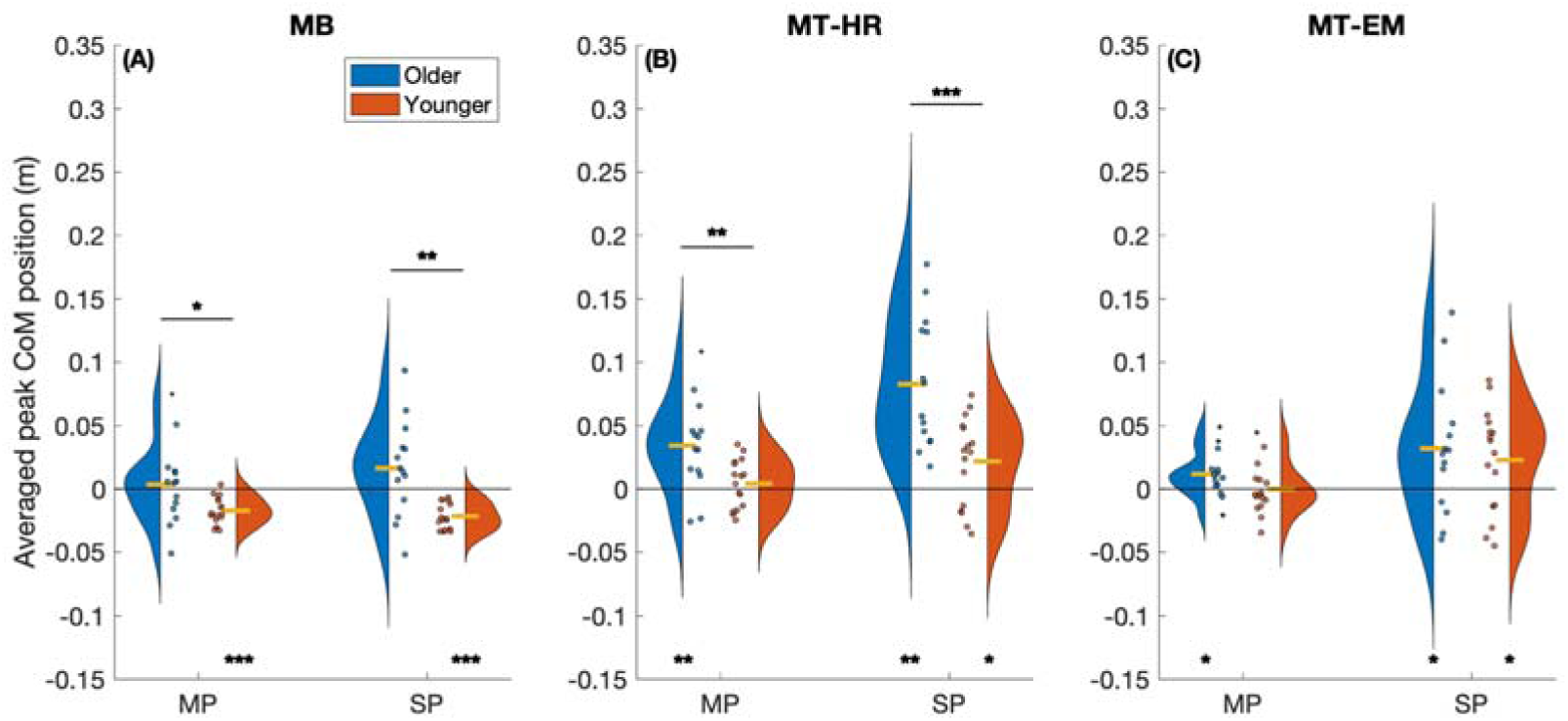
Averaged peak CoM position in both older and younger groups relative to the first sample of the visual perturbation epoch in the moving phase and stationary phase, respectively. The yellow lines represent for the mean values of each group. From left to right: MB: moving background; MT-HR: moving target with head rotation; MT-EM: moving target with eye movement. Significant differences from zero are indicated by an asterisk at the bottom. Significant between groups are indicated by an asterisk on the top

**Table 2.**
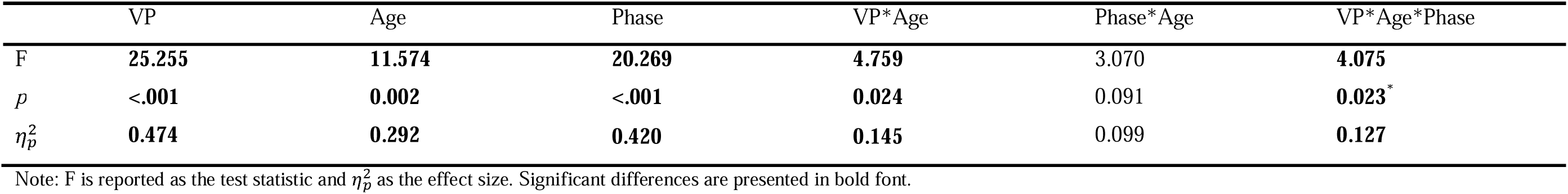
Results of repeated measures ANOVA with three factors (Visual perturbation, Phase and Age) on averaged peak CoM position.

|  | VP | Age | Phase | VP*Age | Phase*Age | VP*Age*Phase |
| --- | --- | --- | --- | --- | --- | --- |
| F | **25.255** | **11.574** | **20.269** | **4.759** | 3.070 | **4.075** |
| *p* | **<.001** | **0.002** | **<.001** | **0.024** | 0.091 | **0.023\*** |
| $\eta_p^2$ | **0.474** | **0.292** | **0.420** | **0.145** | 0.099 | **0.127** |

Note: F is reported as the test statistic and $\eta_p^2$ as the effect size. Significant differences are presented in bold font.

There was a significant three-way interaction effect on averaged peak CoM position. Post-hoc tests indicated that peak CoM position was significantly larger in older adults than in younger adults in both phases in the MT-HR condition (**Table 3**), with values 8.5 and 3.8 times those of the younger adults in the moving and stationary phases, respectively. While in the MB condition, it was smaller in the older adults, which was only 23.5% and 72.7% of the values observed in the moving and stationary phases, respectively. There were no significant differences between older and younger adults in the MT-EM condition.

**Table 3.** Independent t-test results for averaged peak CoM position between older and younger adults during moving and stationary phases in MB, MT-HR and MT-EM conditions.

|  | MB |  |  |  | MT-HR |  |  |  | MT-EM |  |  |  |
| --- | --- | --- | --- | --- | --- | --- | --- | --- | --- | --- | --- | --- |
|  | MP |  | SP |  | MP |  | SP |  | MP |  | SP |  |
|  | older | younger | older | younger | older | younger | older | younger | older | younger | older | younger |
| Mean | **0.004** | **-0.017** | **0.016** | **-0.022** | **0.034** | **0.004** | **0.083** | **0.022** | 0.012 | -1.079*10<sup>-4</sup> | 0.032 | 0.023 |
| 95% CI | **-0.015 –** | **-0.023 –** | **-0.005 –** | **-0.027 –** | **0.014 –** | **-0.006 –** | **0.053 –** | **0.003 –** | 0.001 – | -0.011 – | 0.002 – | 0.001 – |
|  | **0.022** | **-0.011** | **0.038** | **-0.016** | **0.055** | **0.014** | **0.113** | **0.041** | 0.022 | 0.011 | 0.062 | 0.045 |
| Statistic | **-2.317** |  | **-3.661** |  | **-2.908** |  | **-3.861** |  | -1.682 |  | -0.544 |  |
| *p* | **0.035** |  | **0.002** |  | **0.007** |  | **<.001** |  | 0.104 |  | 0.591 |  |
| Cohen’s d | **-0.870** |  | **-1.378** |  | **-1.064** |  | **-1.413** |  | -0.616 |  | -0.199 |  |

Note: MB: moving background; MT-HR: moving target with head rotation; MT-EM: moving target with eye movement. MP: moving phase; SP: stationary phase. Significant differences are presented in bold font.

### The effect of repeated visual perturbation exposure

All statistical results can be found in **Table 4** and **Figure 5**.

**Table 4.** Summary of linear mixed-effects regression analysis of center of mass (CoM) position relative to perturbations numbers in three visual perturbation conditions. Statistical results refer to the older adults. The older group (n = 14) and younger group (n = 16).

|  | Intercept | Repetition | Age | Interaction |
| --- | --- | --- | --- | --- |
| **MB** |  |  |  |  |
| Estimate (95% CI) | **-0.022 (-0.039 - -0.013)** | **0.004 (0.003- 0.005)** | 0.004 (-0.010 - 0.025) | **-0.004 (-0.005 - -0.002)** |
| SE | **0.007** | **0.001** | 0.010 | **0.001** |
| tStat | **-3.115** | **6.361** | 0.363 | **-4.523** |
| pValue | **0.002** | **<.001** | 0.717 | **<.001** |
| **MT-HR** |  |  |  |  |
| Estimate (95% CI) | **0.031 (0.015 - 0.047)** | 0.001 (-0.000 - 0.002) | **-0.028 (-0.049 - -0.006)** | -0.001 (-0.002 - 0.001) |
| SE | **0.008** | 0.001 | **0.011** | 0.001 |
| tStat | **3.869** | 1.400 | **-2.533** | -0.809 |
| pValue | **<.001** | 0.162 | **0.012** | 0.419 |
| **MT-EM** |  |  |  |  |
| Estimate (95% CI) | **0.021 (0.008 - 0.033)** | **-0.002 (-0.003 - -0.000)** | **-0.021 (-0.038 - -0.004)** | **0.002 (0.000 - 0.004)** |
| SE | **0.006** | **0.001** | **0.009** | **0.001** |
| tStat | **3.308** | **-2.433** | **-2.468** | **2.130** |
| pValue | **0.001** | **0.015** | **0.014** | **0.034** |

Note: MB: moving background; MT-HR: moving target with head rotation; MT-EM: moving target with eye movement. Estimate, SE, tStat are referred to the older adults. Significant differences are presented in bold font.

**Figure 5.**
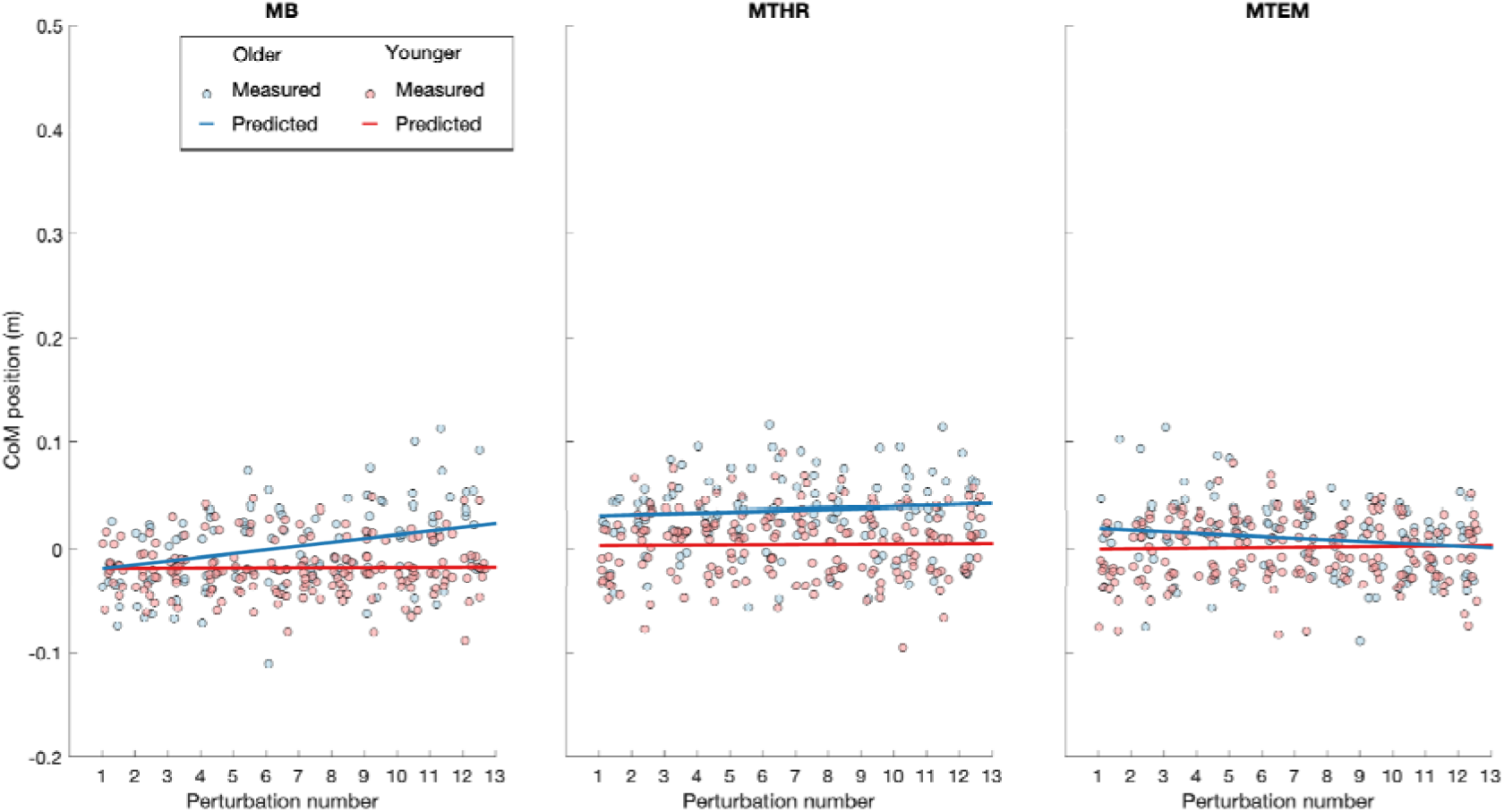
The change of peak CoM position and predicted CoM position by a linear mixed model with visual perturbation repetition numbers during moving phase in three visual perturbation conditions. From left to right: MB: moving background; MT-HR: moving target with head rotation; MT-EM: moving target with eye movement

In MB condition, the CoM position deviated significantly leftward during the first visual perturbation, in both older and younger adults, as indicated by a significant intercept. We found a significant Repetition X Age interaction, with repeated visual perturbations, the CoM position in the older adults shifted from the left to the right side (positive coefficient). In the younger adults, the CoM deviation did not change significantly across perturbation repetitions (i.e. the sum of the Repetition and Interaction coefficients was 0).

In MT-HR condition, the CoM position of the older adults deviated significantly rightward during the first visual perturbation, while the younger adults did not deviate significantly. With visual perturbation repetitions, the CoM position deviation significantly increased in both age groups. There was no significant effect of Repetition or Repetition X Age in the MT-HR condition.

In MT-EM condition, the CoM position deviated significantly rightward during the initial visual perturbations in the older adults, but not in the younger adults. There was a significant interaction effect between age and repetition, the CoM deviation decreased significantly in the older adults, while it did not significantly change across perturbation repetitions in the younger adults.

## Discussion

We assessed effects of visual perturbations on balance control during walking and the changes in these effects over repeated visual perturbations in younger and older adults. We exposed participants to three types of perturbations while they were walking on a treadmill: a moving visual background (MB) and visual tracking of a moving target with either head (MT-HR) or eye movements (MT-EM). CoM position variability increased in both groups due to visual perturbations over the whole trial. During background movements, younger and older adults initially deviated in the direction opposite to the moving background. During visual tracking, both groups deviated in the direction of the target movement. Older adults deviated more than younger adults in the MT-HR condition, with deviations 8.5 and 3.8 times those of younger adults in moving and stationary phases, respectively. While they showed a similar deviation in the MT-EM condition. The two groups responded differently to repeated exposure to visual perturbations. During background movements, the direction of the CoM deviation changed in the older adults only from opposite to background movement to in the direction of background movement. In the MT-HR condition, CoM deviations did not change with repeated exposure in either group. While in the MT-EM condition, CoM deviations did not change with repeated exposure in the younger adults but gradually decreased in the older adults. We will discuss the potential reasons for these findings and their implications with respect to changes in the role of visual information in balance control in different age groups below.

Over the whole trials, both groups showed an increased variability mediolateral CoM position in all visual perturbation conditions, in line with effects of balance perturbations in previous studies [22, 23]. Visual information is integrated with vestibular and other sensory information to estimate self-motion [24, 25]. Changes in such estimates during visual perturbations may cause adjustments in foot placement based on the perceived CoM state, which can in turn perturb the CoM trajectory [26]. Changes in such estimates during visual perturbations may cause adjustments in foot placement based on the perceived CoM state, which can in turn perturb the CoM trajectory [26]. Contrary to our hypothesis, in the current study, the older adults did not show larger increases in CoM position variability during the perturbation conditions (**Figure 2**). One possible explanation for this difference with previous findings [27, 28] is that the visual perturbations in the current study were applied discretely rather than continuously. The period without visual perturbations may have attenuated the age impact of the perturbation on a whole walking trial. In the perturbation epochs, the older adults did exhibit larger CoM deviations.

During visual perturbation epochs in the MB condition, younger adults showed a leftward CoM and foot placement deviation, while deviations in the older adults were consistent with those in the young adults only in the beginning of the experiment. In this condition, when participants fixate on a stationary target, background movement is monitored by peripheral vision. Previous studies found that this may cause an illusion of self-motion in the direction opposite to the background movement [12, 13]. Therefore, the rightward background movement in the current study was expected to cause a perception of leftward self-motion. It seems that the younger adults indeed chose a location for foot placement that extends the base of support in the direction of the perceived movement, to catch the illusory deviation of the CoM. While this has been associated with a subsequent opposite CoM movement in another study [11], we found the CoM deviation to follow this shift in the base of support, suggesting that a sideward translation of the walking path, following the balance correction by shifting foot placement was accepted or not perceived. The older adults responded similar to the younger adults at first, however, they did not exhibit larger CoM deviation than the younger adults. This is inconsistent with the finding of Franz, Francis et al. [28]. However, we found that older adults showed opposite deviations after repeated perturbations (**Figure 5**). This explains why overall smaller CoM deviations were found in the older adults.

There are two possible reasons why the older adults showed different responses over time in the MB condition. One potential reason might be a loss of attention on fixation of the target in the older adults [29]. Possibly older adults started following the background and thus treated it as a foreground, which would lead to an opposite illusion of self-motion. To further test this assumption, we imposed another visual perturbation condition in the same group of participants. In this condition, we performed visual perturbations with simultaneous rightward target and background movements. The participants were instructed to gaze straightforward while walking. A rightward CoM position deviation in the older adults was found, which may support the explanation of a loss of attention on fixation of the target in the older adults (**Figure 6**). However, eye-tracking data would be needed to support this speculative explanation.

**Figure 6.**
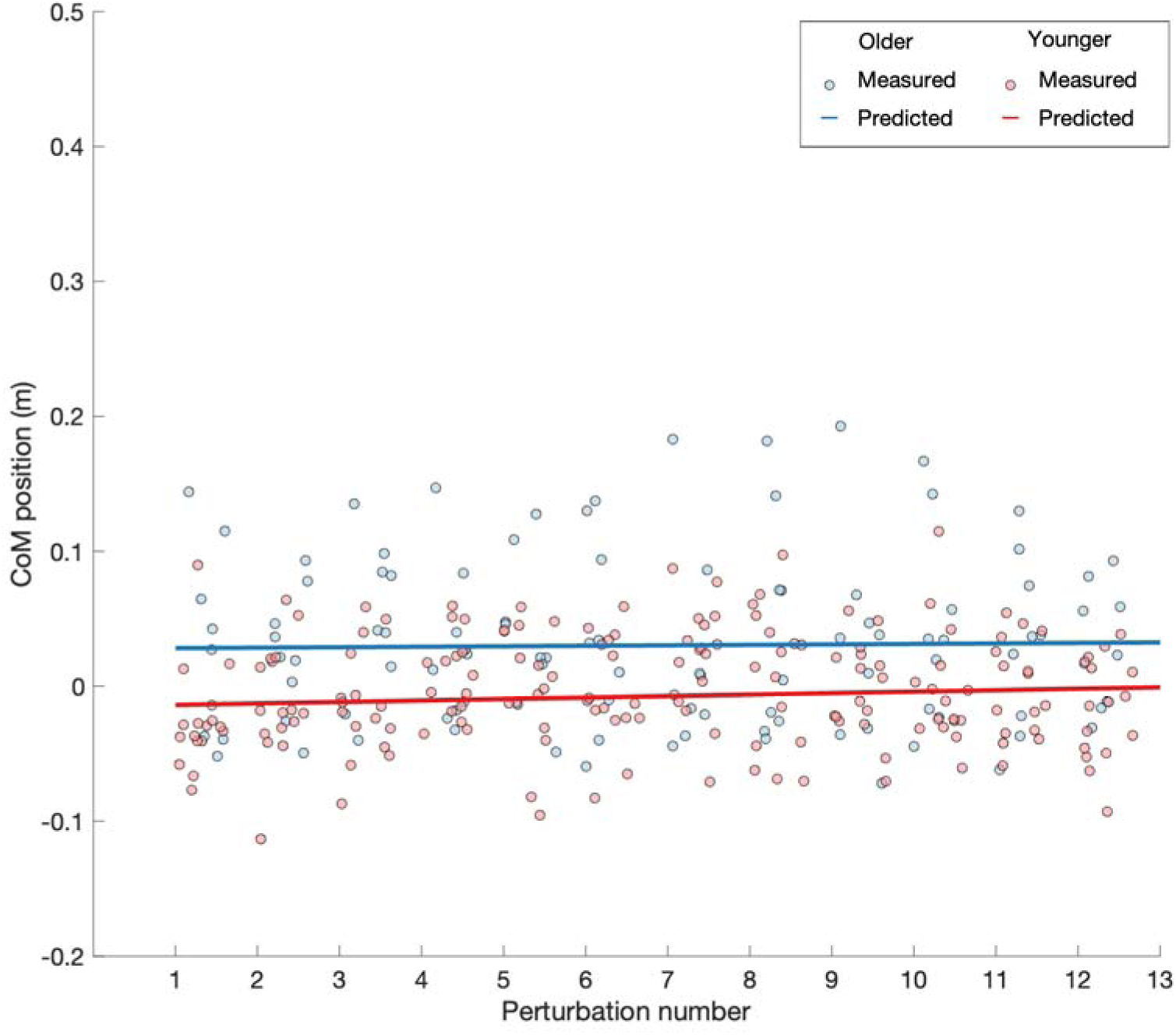
The change of peak CoM position and predicted CoM position by a linear mixed model with visual perturbation repetition numbers during the moving phase in an extra visual perturbation condition. During the visual perturbation, the red target and background moved together to the right while the participants were instructed to maintain a straightforward gaze and head orientation. The result showed a significant intercept only in the older adults, and no significant effect of repetitions during walking

Another reason for the change in deviation direction with repeated exposure in the older adults may be that, after a few perturbations, older adults responded to the perceived self-motion by generating moments in the stance leg to counteract the illusory CoM deviation. This may result in an actual CoM deviation in the direction of the background motion that does not follow the opposite deviation in foot placement demonstrated by Reimann et al. [11, 26], but takes place during the stance phase. This would raise the question why the older adults changed their strategy to respond to an illusory self-motion after repeated exposure. This may be caused by perturbations being harder for the older adults and this becoming harder over time, leading them to choose for the stance leg responses as these take effect more rapidly than foot placement [30].

During visual tracking conditions, tracking the rightward moving target across a stationary background generates a leftward motion of the background images on the retina [31, 32]. The retinal image motion, particularly in the peripheral visual field, contributes to the perception of motion [33]. Thus, in both visual tracking conditions, the rightward movement of the target may produce a leftward motion signal in the peripheral visual field, which could be interpreted as rightward self-motion. The younger adults did not show significant deviations in the moving phases. In contrast, in the older adults, there were substantial rightward foot and CoM deviations from the start of the moving phases. The earlier and stronger responses in the older adults might be a consequence of increased visual dependence. In addition, age-related declines in multisensory integration may contribute to the larger perturbation effects in the older adults. Integration of information from vision, vestibulum, proprioception and efference copies contributes to distinguishing self and externally generated sensations. Wolpe, Ingram et al. [34] found that aging may impair this integration process, thereby diminishing the ability to distinguish between self and externally generated sensations. As a result, older adults may have difficulty accurately interpreting the stationarity of the background during visual perturbations and may be less able to down-regulate perturbation-related visual information compared with younger adults [27, 35].

We found similar peak foot and CoM deviations between the two age groups in the MT-EM condition. However, in the MT-HR condition, the older adults showed larger deviations (**Figure 3 and 4**). A study on rhesus monkeys indicated that the efference copy of the neck motor command gates the processing of vestibular information [36]. Vestibular signals are inhibited during active head rotation where neck proprioceptive information matches the predicted sensory consequences derived from the neck motor command [37]. In line with this, Vallis and Patla [38] found that younger adults preserve heading direction during active head rotation while walking, but not during passive head rotation. Possibly, similar suppression of visual information occurs during target tracking, and this may be more effective in the eye movement only condition, as vestibular and neck proprioceptive information would be unaffected. Less accurate proprioceptive information from the neck in the older adults may lead to failure to suppress self-generated vestibular signals [39], which could explain why the effect of age was more pronounced in MT-HR than in MT-EM.

We expected that downweighting of visual information would occur after several visual perturbations in all conditions, and that the older adults would exhibit a lower ability to downweight the visual inputs. However, the results did not fully agree with our expectations. No consistent repetition effects and interactions of age and repetitions were found. Interactions between age and repetitions on CoM deviations were found only in MB and MT-EM conditions, there were interactions between age and repetitions on CoM deviations. In MB, we surprisingly found that the older adults gradually changed their deviation direction, while the younger adults did not. As we mentioned above, we infer that the change in older adults may be caused by attention distraction or shifting towards faster stance leg responses. In MT-EM, the deviation in the older adults decreased over repetitions, in line with the reweighting theory [40]. The younger adults did not show such a decrease, but their responses were of smaller magnitude to begin with, suggesting that they may have relied more on other sources of sensory information from the beginning. This may suggest that older adults preserve the ability to downweight visual information that is not precise when exposed repeatedly, but that this would not be effective when vestibular and proprioceptive information are also perturbed by head movement. In the MT-HR condition, neither of the groups showed an indication of downweighting of the visual information. This may have been due to the fact that head rotation also affects vestibular and somatosensory information.

Since age-related changes in cognitive and physical status can affect balance control [41, 42], these aspects were measured using clinical scales and were similar in the two age groups. Although a lower score on the DGI test was found in the older adults, all participants were classified as safe ambulators. Still, visual perturbations as applied in the current study were clearly challenging for the older adults. During these conditions, they found it difficult to walk without using the handrail. Three participants deviated excessively from the center of the treadmill and frequently looked down at the ground for additional visual references in both MT-HR and MT-EM conditions, making it necessary to terminate the measurements. This suggests that visual tracking during walking severely challenges older adults, in line with the larger CoM position variability and deviations of CoM trajectories in the target tracking conditions. Two participants could not maintain the speed when walking on the treadmill. Afterwards, they reported that they had fallen in the preceding three months, which they had not reported at inclusion. One participant displayed asymmetric gait when getting fatigued.

### Limitations

The use of a treadmill allowed control of walking speed and collection of data over multiple strides and perturbations per trial. The contact between feet and the moving belts can provide information through proprioceptive and cutaneous input. It may constrain the heading and affect participants’ responses to the visual perturbations.

The high drop-out rate in the older adults may have strengthened a population selection bias that tends to be present in lab studies on older adults in general, i.e. participants in such studies tend to be more fit and active than the general population. While we aimed to recruit healthy older adults with good cognitive status and good balance and gait ability, to minimize any influence of factors other than age, the participants who were able to complete the full experimental protocol are likely to be even more fit and physically active. Therefore, our findings may underestimate effects of ageing in the general population.

For safety reasons, the older adults used a harness. We minimized the potential effect of the harness on their gait by keeping the harness slack, so that tension would be applied only when the participants required support to prevent a fall. However, the harness may have provided some sensory cues when they deviated from the center of the treadmill. In that case, we will have underestimated the effects of age.

We designed our experiment and interpreted results based on effects of visual information on balance control. Visual information together with head rotation or gaze direction plays a role in control of heading, and effects on heading and balance are not easy to distinguish. Visual perturbation used in current study may have influenced balance control as well as heading direction. However, we are not able to disentangle the effects of visual perturbations on control of balance from those on heading. Future work will investigate the possible way to distinguish the effect of visual information on heading and balance control.

In the current study, downweighting on visual information following repeated visual perturbations was observed only in the MT-EM condition. The duration of the perturbations may not have been sufficient to elicit reweighting in the other conditions.

### Conclusions

The present findings showed that the older adults had lower ability to control CoM trajectories in the frontal plane when exposed to horizontal visual perturbations during walking. This appears to be driven by impaired self-motion estimation with aging. Older adults exhibit lesser ability to accurately estimate self-motion through correction by other sensory sources when exposed to visual perturbations. However, they still maintain the capacity to downweight visual information after repeated exposure to visual perturbations.

## Acknowledgements

This study was supported by a scholarship (No. 202308310074) from the China Scholarship Council (CSC). We appreciate all participants for their voluntary participation. We thank Leon Schutte, Richard Casius, and Paul Pesman for the technical assistance. We also thank Robbert Verkuil, Raven Huiberts and other collogues for their help in data collection.

## Data and resource availability

All data have been made available online. All files can be downloaded from: https://doi.org/10.5281/zenodo.20141231

