## Supplemental material for "Balance control is more affected in older than younger adults by repeated visual perturbations during walking"

**Gait characteristics over the whole walking trial**

Here we provide additional information on gait characteristics over the whole walking trial including step width, step width variability, and step frequency (**Figure S1**).

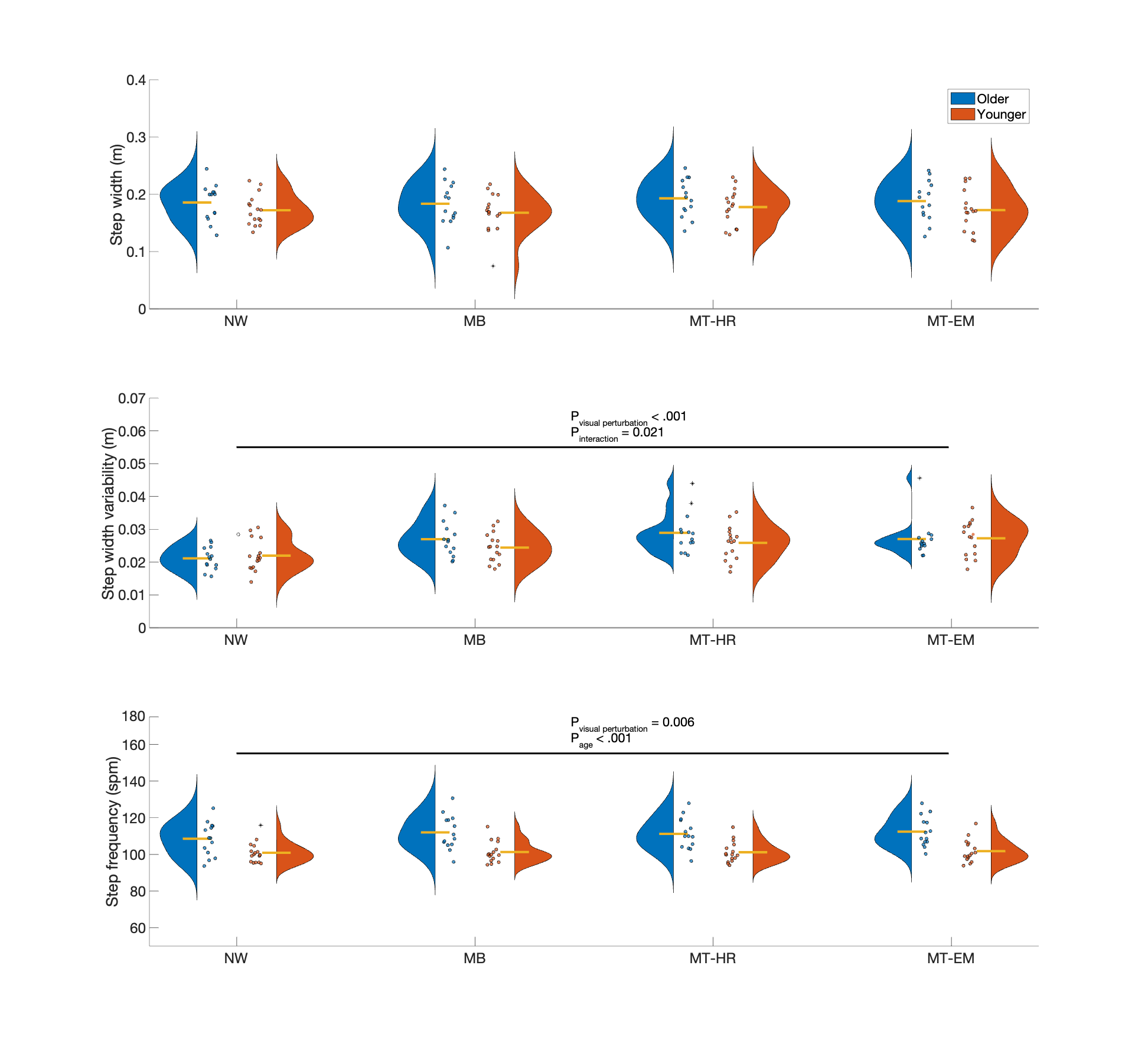

Figure S1 Effects of Visual perturbation conditions on step width, step width variability and step frequency in older and younger adults. The yellow lines represent for the mean values of each group. NW: normal walking, MB: moving background, MT-HR: moving target with head rotation, MT-EM: moving target with eye movement.

There was no significant main effect of age or visual perturbation, and their interaction on step width over the whole walking trial.

There was a significant interaction between Age and Visual perturbation on the step width variability (F _age*visual perturbation_ (df = 3, 84) = 3.410, *p* = 0.021, $\eta_{p}^{2}$= 0.109), indicating that the effect of visual perturbations differed between age groups. Whereas there was no significant main effect of age. Compared with normal walking, the step width variability in the older adults significantly increased in the MB (*p* < .001, *d* = 1.156), MT-HR (*p* < .001, *d* = 1.537) and MT-EM (*p* = 0.002, *d* = 1.161) conditions.

There was no significant interaction effect on step frequency. The older adults showed larger step frequency than the younger adults (F _age_ (df = 1, 28) = 14.187, *p* < .001, $\eta_{p}^{2}$= 0.336; post hoc results: *p* < .001, *d* = 1.327).

**Averaged peak foot position in one visual perturbation epoch**

In the MB condition, in the older participants, averaged peak foot positions were not significantly different from zero in the moving or stationary phase (*p* = 0.756, *d =* 0.085; *p* = 0.165, *d =* 0.393), in contrast with the younger participants who showed significant negative deviations in both phases (*p* <.001, *d* = -1.663; *p* <.001, *d* = -1.861). In the MT-HR condition, in older adults, peak foot positions were significantly larger than zero in both moving and stationary phases (*p* = 0.002, *d* = 1.065; *p* < .001, *d* = 1.673), indicating positive deviations in the target direction in both phases. The younger participants only showed significant positive deviations in the stationary phase (*p* = 0.018, *d* = 0.662). In the MT-EM condition, foot position deviations were not significantly different from zero in the moving phase in both age groups (older: *p* = 0.105, *d* = 0.466; younger: *p* = 0.827, *d* = -0.056). In the stationary phase, the older adults significantly deviated in the direction of the target (*p* = 0.040, *d* = 0.610), as did the younger participants (stationary phase: *p* = 0.039, *d* = 0.565).

There was a significant Visual perturbations X Age interaction effect on peak foot positions. Post-hoc tests showed that peak foot positions were larger in the older participants in MB (*p* = 0.035, *d =* 0.902) and MT-HR conditions (*p* = 0.006, *d =* 1.433).

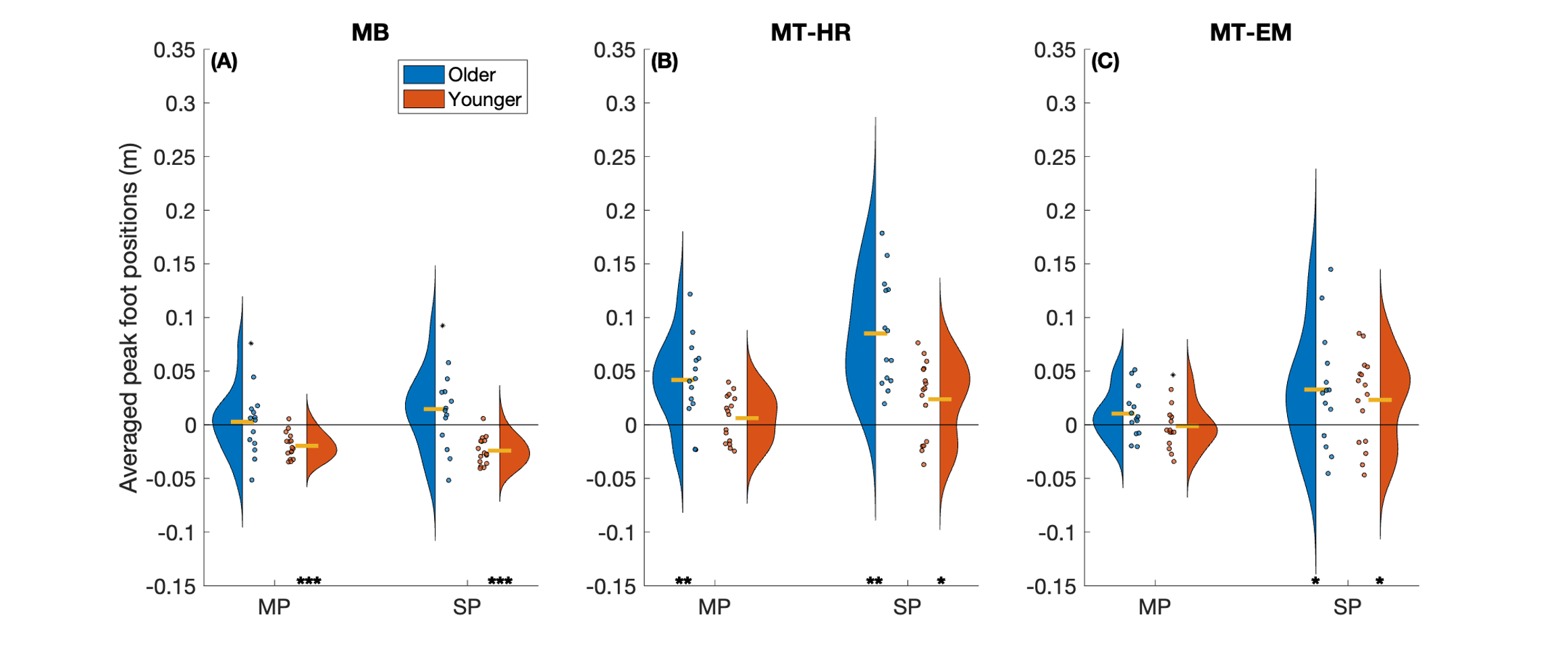

Figure S2 Averaged peak foot positions in both older and younger groups relative to the first sample of the visual perturbation epoch in the moving phase and stationary phase, respectively. The yellow lines represent for the mean values of each group. From left to right: MB: moving background; MT-HR: moving target with head rotation; MT-EM: moving target with eye movement. Significant differences from zero are indicated by an asterisk.

Table 1 Results of repeated measures ANOVA with three factors (Visual perturbation, Phase and Age) on averaged peak foot positions.

|  | **VP** | **Age** | **Phase** | **VP*Age** | **Phase*Age** | **VP*Age*Phase** |
| --- | --- | --- | --- | --- | --- | --- |
| **F** | **30.397** | **11.543** | **20.678** | **5.175** | 2.503 | 2.751 |
| ***p*** | **<.001** | **0.002** | **<.001** | **0.018** | 0.125 | 0.076 |
| $\boldsymbol{\eta}_{\boldsymbol{p}}^{\boldsymbol{2}}$ | **0.521** | **0.292** | **0.425** | **0.156** | 0.082 | 0.089 |

Note: F is reported as the test statistic and $\eta_{p}^{2}$ as the effect size. Significant differences are presented in bold font.
